# A global survey of intramolecular isopeptide bonds

**DOI:** 10.1101/2025.04.25.650565

**Authors:** Francesco Costa, Ioannis Riziotis, Antonina Andreeva, Delhi Kalwan, Jennifer de Jong, Philip Hinchliffe, Fabio Parmeggiani, Paul R. Race, Steven G. Burston, Alex Bateman, Rob Barringer

**Affiliations:** European Molecular Biology Laboratory, European Bioinformatics Institute (EMBL-EBI), Wellcome Genome Campus, Hinxton. CB10 1SD. UK; School of Biochemistry, University of Bristol, University Walk, Bristol. BS8 1TD. UK; School of Cellular and Molecular Medicine, University of Bristol, University Walk, Bristol, BS8 1TD, UK; School of Chemistry, University of Bristol, Cantock’s Close, Bristol. BS8 1TS. UK; School of Pharmacy and Pharmaceutical Sciences, Cardiff University, Redwood Building, Cardiff. CF10 3NB, UK; School of Natural and Environmental Sciences, Devonshire Building, Newcastle University, Newcastle upon Tyne. NE1 7RU. UK

**Keywords:** Fibrillar adhesin, Pilus, Biofilm, Host-Pathogen, Adhesion, Isopeptide

## Abstract

Many protein domains harbour covalent intramolecular bonds that enhance their stability and resistance to thermal, mechanical and proteolytic insults. Intramolecular isopeptide bonds represent one such covalent interaction, yet their distribution across protein domains and organisms has been largely unexplored. Here, we sought to address this by employing a large-scale prediction of intramolecular isopeptide bonds in the AlphaFold database using the structural template-based software Isopeptor. Our findings reveal an extensive phyletic distribution in surface proteins resembling fibrillar adhesins and pilins. All identified intramolecular isopeptide bonds are found in two structurally distinct folds, CnaA-like or CnaB-like, from a relatively small set of related Pfam families, including ten novel families that we predict to contain intramolecular isopeptide bonds. One CnaA-like domain of unknown function, DUF11 (renamed here to “CLIPPER”) is broadly distributed in cell-surface proteins from Gram-positive bacteria, Gram-negative bacteria, and archaea, and is structurally and biophysically characterised in this work. Using X-ray crystallography, we resolve a CLIPPER domain from a Gram-negative fibrillar adhesin that contains an intramolecular isopeptide bond and further demonstrate that it imparts thermostability and resistance to proteolysis. Our findings demonstrate the extensive distribution of intramolecular isopeptide bond-containing protein domains in nature, and structurally resolve the previously cryptic CLIPPER domain.

## Introduction

The isopeptide bond is a class of covalent amide bond that forms between the amino and carboxyl groups of two side chains of a polypeptide or between a side chain and a terminus. In nature, isopeptide bonds enable either the cross-linking of two points within a single polypeptide chain (intramolecular isopeptide bonds) or between two points of two different polypeptide chains (intermolecular isopeptide bonds). Enzyme-catalysed intermolecular isopeptide bonds play a role in many biological processes (Kang and Baker, 2011), including protein ubiquitination, where the target protein is covalently tethered to the ubiquitin protein (Hershko and Ciechanover, 1998); the coagulation pathway, where Factor XIII catalyses crosslinking of fibrin (Muszbek, Adany & Mikkola, 1996); and in pilus assembly, whereby pilins are crosslinked to the pentaglycine peptidoglycan bridge at the microbial cell surface (Hendrickx et al. 2011). Notably, some intermolecular isopeptide bonds form through an autocatalytic mechanism, such as between capsid subunits of various bacteriophages (Helgstrand et al, 2003; Podgorski et al. 2023), where the isopeptide bonds form within a hydrophobic pocket between lysine and asparagine side chains, and are catalysed by a nearby glutamate side chain (Tso et al, 2017).

Over the past two decades a second subclass of autocatalytic isopeptide bond has been investigated, the intramolecular isopeptide bond. The first intramolecular isopeptide bond to be structurally resolved was by Kang et al. in 2007, housed within the hydrophobic core of a β-sandwich domain of the bacterial pilin Spy0128. This isopeptide bond was formed between the lysine and asparagine (Lys-Asn) side chains of adjacent β-strands, catalysed by a proximal glutamate side chain (Kang et al. 2007). Subsequently, several other intramolecular isopeptide bond domains have been identified, invariably formed between side chains within the core of β-sandwich folds. These folds can be grouped into two distinct structural families sharing a Greek-key motif, CnaA-like domains (an Ig-like fold that typically forms Lys-Asn cross-links catalysed by an aspartate) and CnaB-like domains (a transthyretin-like fold that typically forms Lys-Asn or Lys-Asp cross-links catalysed by a glutamate, Kang & Baker, 2009; Kang & Baker, 2012). In CnaA-like domains, intramolecular isopeptide bond-forming residues are located on opposing β-sheets between the first and penultimate β-strands, while in CnaB-like domains they are positioned on the same β-sheet, between adjacent first and last β-strands. Interestingly, both folds are tolerant of domain insertions within loop regions, usually of intramolecular isopeptide or adhesion domains (Izoré et al. 2010; Pointon et al., 2010; Spraggon et al. 2010; Figure 1).

**Figure 1.**
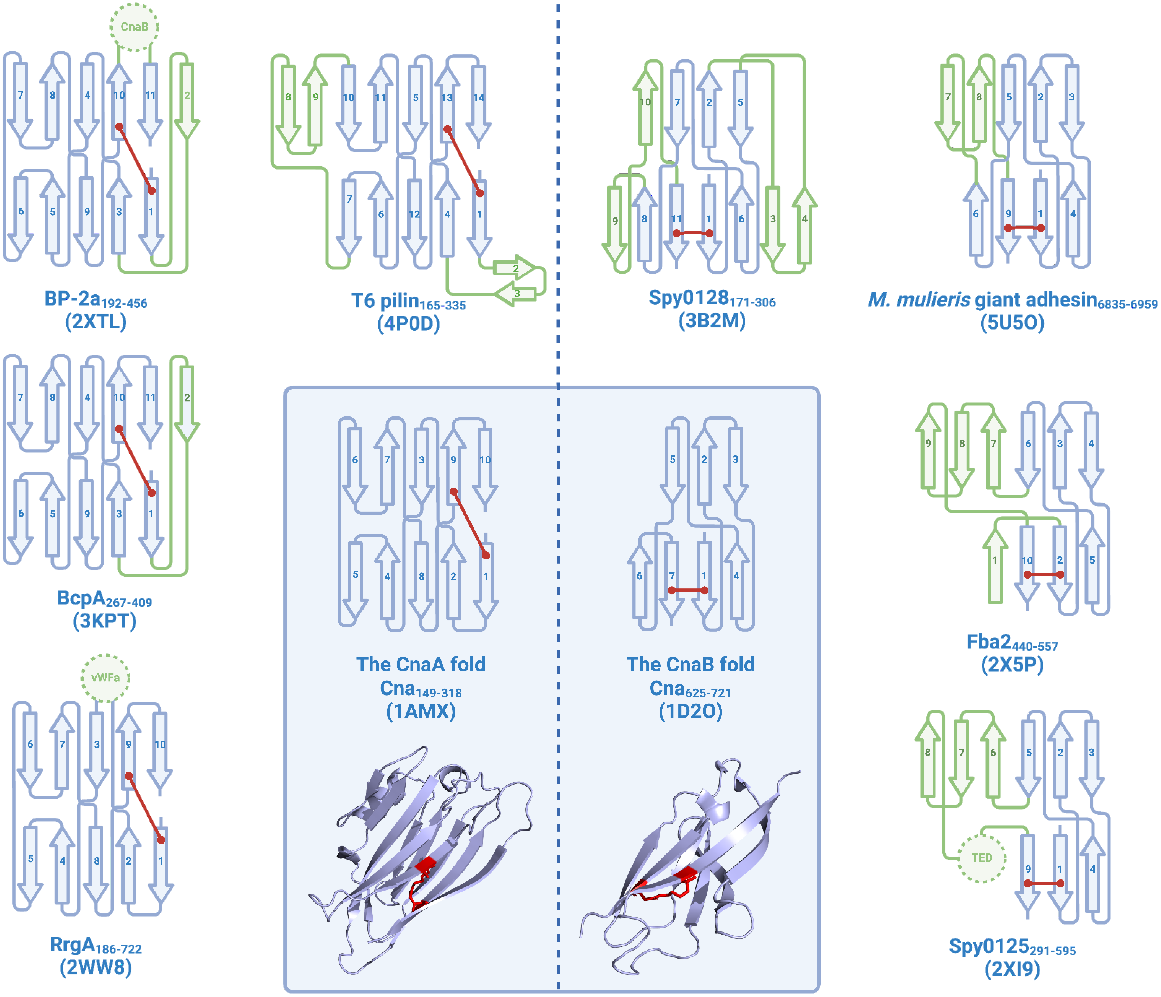
β-strand arrangement in CnaA and CnaB folds. Isopeptide bonds are indicated by red lines, along with cartoon depictions of the folds. Alternative CnaA/CnaB topologies are depicted in the periphery. Differences relative to archetypal CnaA/CnaB folds are highlighted in green. Domain inserts are depicted as dotted circles and labelled as follows: CnaB, CnaB-like fold; vWFa, von Willebrand factor type A domain; TED, Thioester domain.

Functionally, intramolecular isopeptide bonds are thought to enable resistance to various stresses at the cell surface. In 2007, Kang et al. demonstrated that intramolecular isopeptide bonds bestow increased proteolytic resistance and thermostability to the CnaB-like domains of Spy0128, which later proved characteristic of CnaA-like and CnaB-like domains more broadly (Kang et al. 2007; Kang & Baker, 2009; Kang & Baker, 2011; Zähner et al., 2011; El Mortaji et al. 2012; Hendrickx et al. 2012; Chaurasia et al., 2016; Heidler et al. 2021). Further work revealed that the intramolecular isopeptide bonds of CnaB-like domains in Spy0128 enable resistance to mechanical unfolding, with these polypeptides proving inextensible under atomic force microscopy (AFM)-induced tension (Alegre-Cebollada et al. 2010). In contrast, CnaA-like folds were found to unfold partially under tension, leading to the molecular “shock absorber” hypothesis whereby CnaA-like domains enable adherence under shear forces by dissipating mechanical perturbations (Echelman et al. 2016).

To date, intramolecular isopeptide bonds have primarily been identified in adhesive pili and fibrillar adhesins of Gram-positive bacteria, leading some to question whether they may also be found in Gram-negative bacteria, archaea, eukaryotes or viruses (Schwarz-Linek & Banfield, 2014). This open question was partially answered in 2021, when an intramolecular isopeptide bond was identified in a pilin of a Gram-negative bacteria (Heidler et al. 2021). Despite this finding, the prevalence and phyletic distribution of intramolecular isopeptide bond domains in nature has not been systematically probed, and the proteins harbouring these domains have not been systematically characterised.

Here, we present the first large-scale prediction of naturally occurring intramolecular isopeptide bonds in the AlphaFold Database, providing insights into their distribution across organisms and protein domains. Using X-ray crystallography, we subsequently resolve an intramolecular isopeptide bond in DUF11, a domain of unknown function found in fibrillar adhesins of Gram-positive bacteria, Gram-negative bacteria, and archaea, and confirm that the isopeptide bond imparts significant thermostability and proteolytic resistance. Our work reveals that intramolecular isopeptide bonds frequently appear in stalks of bacterial and archaeal fibrillar adhesins, and likely facilitate adherence under various stressful conditions.

## Results

### Characteristics of intramolecular isopeptide bonds

Previous experimental and computational studies revealed that hydrophobic environments likely facilitate intramolecular isopeptide bond formation (Hagan et al. 2010; Hu et al. 2011). However, a comprehensive analysis of the environment of intramolecular isopeptide bonds has not yet been undertaken. Consequently, we first quantified the solvent accessibility of intramolecular isopeptide bonds within protein structures deposited in the PDB, and characterised the physicochemical properties of the environment surrounding the bond.

Using a PDB dataset of all known intramolecular isopeptide structures collated in prior work (Costa et al. 2025), we found that the relative solvent accessible surface area (rASA) of most intramolecular isopeptide bonds was <0.05, confirming the buried nature of isopeptide bonds within hydrophobic cores (Kang et al. 2007) (Figure 2A). When assessing *cis*/*trans* conformations of intramolecular isopeptide bonds, we found that no Lys-Asp bonds are present in the *cis* conformation, while Lys-Asn bonds are equally found in either *cis* (50%) or *trans* (48%) conformations (with 2% found in intermediate conformations). This builds on our previous observation that 67% of CnaB-like domains favour the *cis* conformation while 73% of CnaA-like domains favour the *trans* conformation (Costa et al. 2025). We also quantified the frequency of proximal aromatic side chains (a feature which has only been qualitatively observed to date, Kang et al. 2007; Kang & Baker, 2011), and noted that the Nζ atom of the isopeptide bond is often located 3-6 Å above the plane of an aromatic sidechain (Figures 2B and 2D). These “aromatic caps” (as we term them here) appear more prevalent in CnaA-like domains (Figure 2C) but are notably absent in folds harbouring insertions of large domains between isopeptide bond-contributing residues (e.g. in the the CnaA-like domain of the RrgA pilin and the CnaB-like domain of the Spy0125 pilin, Izoré et al. 2010; Pointon et al. 2010). Aromatic caps are more prevalent in *cis* Lys-Asn intramolecular isopeptide bonds (86% vs 29% prevalence in *cis*/*trans* conformations, respectively), and are absent in all PDB structures containing Lys-Asp bonds. Our analyses also frequently revealed the presence of a water molecule within 5 Å of the intramolecular isopeptide bond oxygen in 54% of CnaB-like and 59% of CnaA-like domains, usually buried in the domain core.

**Figure 2.**
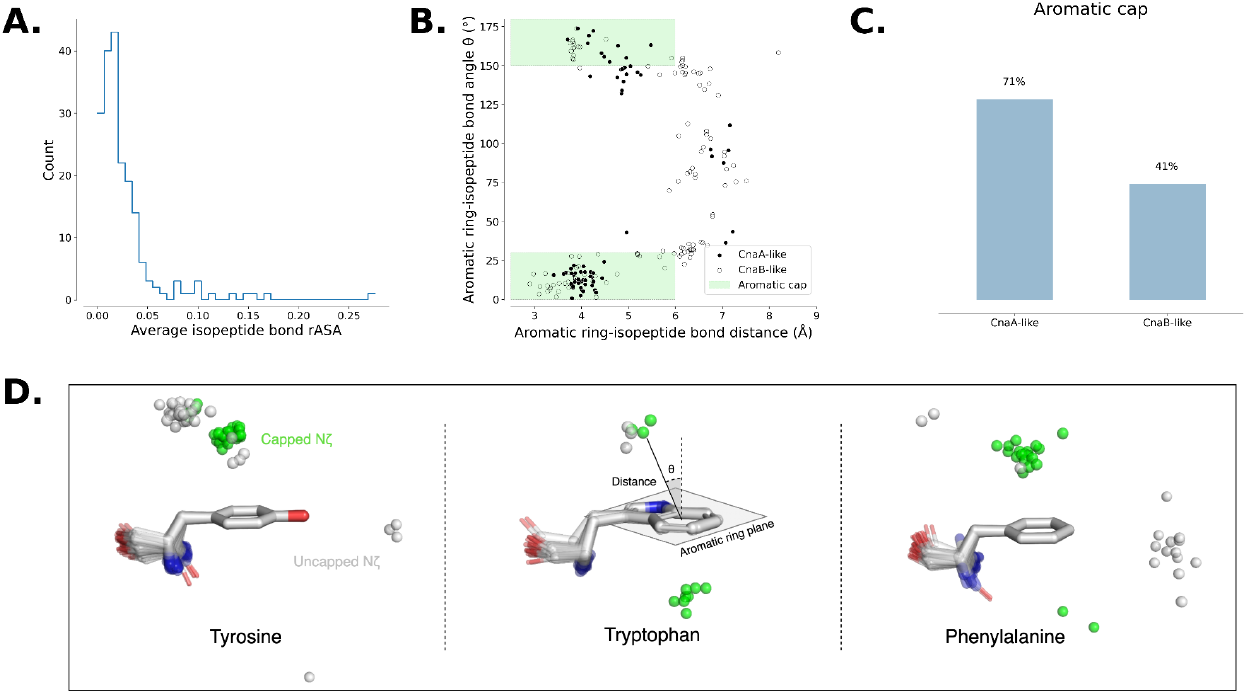
A. Relative accessible solvent area (rASA) averaged across the three isopeptide bond residues. B. Scatter plot displaying distance and angles between the intramolecular isopeptide bond and aromatic ring plane (distance calculated between the lysine Nζ atom and the centroid of the aromatic ring, angle calculated between the lysine Nζ atom-centroid vector and the normal of the aromatic ring plane). C. Percentage of intramolecular isopeptide bonds with an aromatic cap, defined as an isopeptide bond within 6 Å of the aromatic ring plane, at an angle <30**°** or >150**°** from the normal of the aromatic ring plane (see panel B). Only one PDB entry was assessed per sequence-identical domain. D. Positioning of Nζ isopeptide bond atoms around their proximal aromatic ring. “Capped Nζ” refers to isopeptide bonds demonstrating an aromatic cap relationship with a proximal aromatic ring.

### Large-scale prediction of intramolecular isopeptide bonds in the AlphaFold Database

We next employed a structure-guided search method to identify putative intramolecular isopeptide bonds within predicted structures in the AlphaFold Database (AFDB, Varadi et al. 2024) using the isopeptide-scanning software Isopeptor, which identifies isopeptide bonds in structural models using a template-based matching approach and assigns probability scores to each hit (Costa et al. 2025). We initially tested Isopeptor’s ability to detect intramolecular isopeptide bonds in AlphaFold2 (AF2) models of PDB-deposited intramolecular isopeptide bond structures to confirm that AF2 reliably approximates the positions of isopeptide bond residues. Isopeptor identified intramolecular isopeptide bonds in 94% of AF2 models using a probability threshold of 0.65 (comparable to the recall in PDB structures, Costa et al. 2025), confirming that AF2 places intramolecular isopeptide bond residues in positions consistent with those found in PDB depositions (Figure S1).

Applying Isopeptor to the AFDB using a probability threshold of 0.65, we identified 69,718 intramolecular isopeptide bonds within 33,049 (0.015%) of the 214,683,839 models, all of which were located within β-sandwich folds. Since the AFDB does not contain predictions of viral proteins, a scan was also performed against the Big Fantastic Virus Database (BFVD) to determine whether intramolecular isopeptide bonds could be detected in viral proteins (Kim et al. 2025). No hits were returned from the BFVD (data not shown). To characterise the identified intramolecular isopeptide bond domains into distinct families, we mapped the AFDB Isopeptor hits to Pfam protein domains. The identified domains fell into 26 Pfam families, 12 of which had been structurally characterised with an intramolecular isopeptide bond (Figures 3A and S2, Tables 1 and S2). Multiple hits matched to four pre-existing families, two that had been annotated as probable intramolecular isopeptide bond domains (DUF11 and SpaA_3), and two that had not (SdrD_B and SpaA_2). Many hits did not belong to existing Pfam families and were, therefore, used as seeds to create ten new Pfam families. All 26 families can be grouped into three superfamilies according to the Pfam classification: the Adhesin (CL0204), Transthyretin (CL0287) and E-set (CL0159) clans.

**Table 1.** Pfam domains confirmed by structural characterisation (bold) or predicted (plain) to harbour intramolecular isopeptide bonds.

| Pfam ID<br>Pfam (accession code) | Pfam Clan ID<br>(accession code)-isopeptide bond class type | PDB/AFDB Accession Code | PDB chain | Pfam Domain start-end | Structural Domain start-end | Experimental/predicted intramolecular isopeptide bond residues (first bond residue, catalytic residue, second bond residue) |
| --- | --- | --- | --- | --- | --- | --- |
| **AgI_II_C2 (PF17998)** | Adhesin (CL0204)<br>CnaA-like | 5DZ9 | A | 554-723 | 554-723 | 556, 606, 703 |
| **Sgo0707_N2 (PF20623)** |  | 4HSS | A | 178-324 | 184-324 | 187, 224, 299 |
| **GramPos_pilinBB (PF16569)** |  | 2X9Z | A | 216-328 | 187-325, 436-446 | 193, 241, 318 |
| **Collagen_bind (PF05737)** |  | 7LGR | A | 173-306 | 173-317 | 177, 210, 291 |
| **Antigen_C (PF16364)** |  | 3OPU | A | 1334-1492 | 1330-1489 | 1334, 1383, 1469 |
| DUF7926 (PF25548) |  | A0A179B332 | - | 188-359 | 190-349 | 193, 241, 334 |
| DUF7929 (PF25551) |  | A0A090G7K7 | - | 81-255 | 74-257 | 86, 129, 213 |
| DUF7925 (PF25546) |  | Q8YS00 | - | 259-442 | 260-508 | 264, 326, 415 |
| **DUF5979 (PF19407)** | Transthyretin (CL0287)<br>CnaB-like | 4BUG | A | 606-718 | 603-718 | 610, 680, 715 |
| **SpaA (PF17802)** |  | 2XID | A | 291-377 | 291-384, 588-598 | 297, 347, 595 |
| **GramPos_pilinD3 (PF16570)** |  | 2X9Z | A | 388-448 | 344-433 | 349, 405, 428 |
| **FctA (PF12892)** |  | 7W7I | A | 48-216 | 37-218 | 52, 139, 213 |
| **GramPos_pilinD1 (PF16555)** |  | 7WOI | A | 48-196 | 48-196 | 57, 158, 195 |
| **Cna_B (PF05738)** |  | 1D2O | A | 627-716 | 625-721 | 633, 694, 717 |
| SpaA_2 (PF19403) |  | A0A086ZQG5 | - | 489-625 | 489-629 | 497, 571, 627 |
| SpaA_3 (PF20674) |  | A0A514BTT6 | - | 293-411 | 291-412 | 298, 353, 409 |
| SpaA_4 (PF24514) |  | A0A4R4IAB0 | - | 700-816 | 696-817 | 704, 766, 814 |
| SdrD_B (PF17210) |  | A0A7C3F8X4 | - | 157-243 | 153-250 | 159, 218, 248 |
| DUF7601 (PF24547) |  | B8QYD3 | - | 184-307 | 184-308 | 190, 263, 305 |
| **GBS104-like_lg (PF21426)** | E-set (CL0159)<br>CnaA-like | 3TXA | A | 587-717 | 137-211, 587-717 | 188, 597, 692 |
| DUF11 (PF01345) |  | 9IFR (from this work) | A | 712-812 | 708-828 | 715, 748, 806 |
| DUF7507 (PF24346) |  | A0A2I2KYV6 | - | 65-169 | 65-176 | 72, 112, 152 |
| DUF7619 (PF24595) |  | L7WFS8 | - | 653-784 | 655-785 | 657, 695, 764 |
| DUF7927 (PF25549) |  | A0A6G8FG97 | - | 120-250 | 119-242 | 125, 161, 225 |
| DUF7617 (PF24593) |  | A0A495X870 | - | 676-804 | 678-807 | 684, 740, 778 |
| DUF7933 (PF25564) |  | A0A023BX92 | - | 968-1093 | 968-1096 | 973, 1008, 1068 |

**Figure 3.**
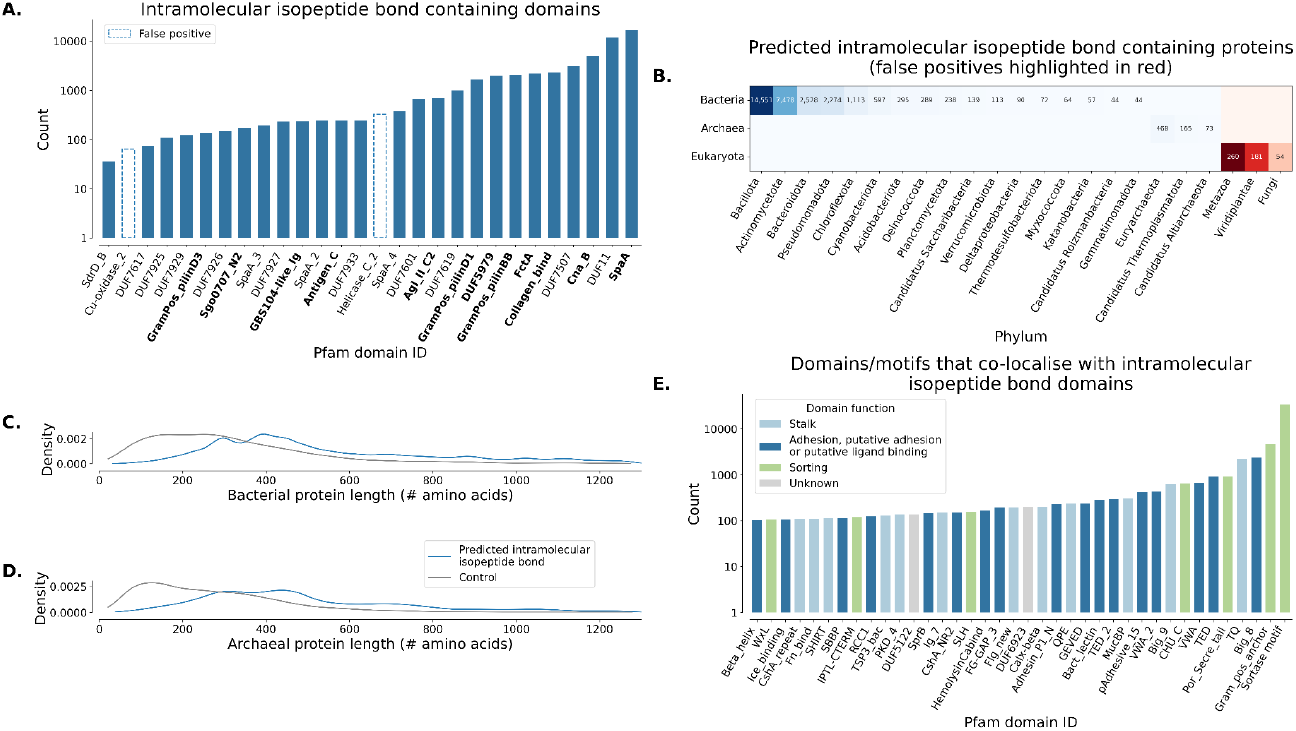
A. Pfam domains predicted by Isopeptor to contain intramolecular isopeptide bonds in the AFDB (probability >0.65). Domains in bold were structurally characterised with an intramolecular isopeptide bond prior to this work. B. Life domain and phylum distribution of intramolecular isopeptide bond-containing proteins detected with Isopeptor in the AFDB (probability >0.65, phyla with <40 sequences are not shown). Eukaryotic proteins are likely false positives or are from incorrectly annotated organisms. C/D. Sequence length distribution of bacterial and archaeal intramolecular isopeptide bond proteins detected by Isopeptor in the AFDB. Length distributions are compared with a set of 10,000 randomly selected AFDB proteins from each life domain. Note that the AFDB has an upper sequence length limit of 1,280 amino acids. E. Pfam domains neighbouring isopeptide bond domains identified by Isopeptor in AFDB proteins. Domains detected <100 times are not shown (full data are available in Figure S5), nor are domains from polypeptides annotated as eukaryotic.

With an updated collection of intramolecular isopeptide bond domain families, we then surveyed their phyletic distribution in the AFDB. This revealed that intramolecular isopeptide bond domains are predominantly found in bacterial entries, but we also identified hits within archaea and eukaryotes (Figure 3B). While most intramolecular isopeptide domains are found within the Gram-positive Bacillota and Actinomycetota phyla, a significant number are found within the Gram-negative Bacteroidota and Pseudomonadota phyla. Two previously uncharacterised DUFs were identified in Gram-positive bacteria, Gram-negative bacteria, and archaea: DUF11 and DUF7507, with DUF11 being by far the most broadly distributed (Figure S3). While hits were identified in eukaryotes, manual inspection revealed them to be likely false positives as the residues were solvent-exposed, not conserved across families, and lacked homologous proteins in related eukaryotes, suggesting that these sequences may have originated from microbial contamination during sequencing projects.

We noted that the proteins identified by Isopeptor are typically longer than other proteins and often harbour tandemly repeating intramolecular isopeptide domains (Figures 3C, 3D and S4). These domains are often neighboured by adhesive domains, particularly Big_8 (a putative adhesion domain related to the collagen-binding domain of the adhesin CNA, Zong et al. 2005) and also stalk domains such as the threonine-glutamine domain (TQ), which possesses a stabilising intramolecular ester bond (Kwon et al. 2014). Other common neighbouring domains include sorting domains such as the Por_secre_tail, the CHU_C domain, and the LPXTG cell wall anchor, which enable anchoring to the cell surfaces of Gram-negative and Gram-positive bacteria, respectively (Figures 3E and S5). Taken together, these data suggest that a sizable portion of these intramolecular isopeptide bonds occur in elongated surface-anchored monomeric proteins that enable adhesion, also known as fibrillar adhesins. Subsequent analysis performed using software for fibrillar adhesin detection (Monzon & Bateman, 2022) revealed that 14% of intramolecular isopeptide bond-containing proteins identified in the AFDB by Isopeptor are likely to be fibrillar adhesins. Indeed, inspection of the Isopeptor hits identified several putative fibrillar adhesins in Gram-positive bacteria, Gram-negative bacteria and archaea, covering pathogens, opportunistic pathogens, and commensal microbes (Figure S6, Table S3).

Fibrillar adhesins typically present as domain-shuffled polypeptides with unique functionalities, dictated by the specific domain arrangements within the protein structure (Monzon, Lafita & Bateman, 2021; Barringer et al. 2023). Thus, we next characterised neighbouring domains of intramolecular isopeptide bond domains in AFDB polypeptides. Our analyses found a diverse set of neighbouring domains which can be broadly divided into three functional categories: adhesion, stalk-forming and sorting domains (Figure 3E), mainly occurring in bacteria (Figure S5). While intramolecular isopeptide bond domains (primarily DUF11) were identified in archaeal AFDB proteins, neighbouring domains were rarely detected, indicating that either these structures contain only DUF11 repeats or that neighbouring domains do not resemble the current collection of known Pfam domain families. When other domains are present in archaea, they commonly include the Chlam_PMP domain (Pfam ID PF02415), which is also found in Chlamydia surface proteins, and PKD_4 domains (Pfam ID PF18911), which have also been detected in bacterial surface proteins.

### DUF11 is an intramolecular isopeptide bond-containing domain that is widely distributed in fibrillar adhesins of bacteria and archaea

The DUF11 domain family was initially deposited in the Pfam database in 1998 (Pfam ID: PF01345) and was rebuilt in 2018 to better represent several immunoglobulin-like domains, some of which were predicted to engage in intramolecular isopeptide bond formation. AF2 predicts that DUF11 domains fold into an Ig-like β-sandwich structure containing a Greek-key motif and consisting of seven to nine β-strands. While DUF11 belongs to the E-set Ig-like fold clan (CL0159), it shares some topological similarity to domains of the bacterial adhesin clan (CL0204), which both represent CnaA-like domains (Table 1). Typically, DUF11 domains are found in one or in multiple copies, often tandemly repeated in the middle of long proteins harbouring a signal peptide, adhesion domains and sorting domains, indicating that the domain routinely forms the stalk of fibrillar adhesins (Figure 4). While all DUF11 domains share global sequence similarity and are predicted to exhibit a CnaA-like fold, some DUF11 domains appear to lack intramolecular isopeptide bonds (Table S2). This heterogeneity is evident from the Pfam SEED and FULL alignments, in which the positions of the residues predicted to form isopeptide bonds are not universally conserved.

**Figure 4.**
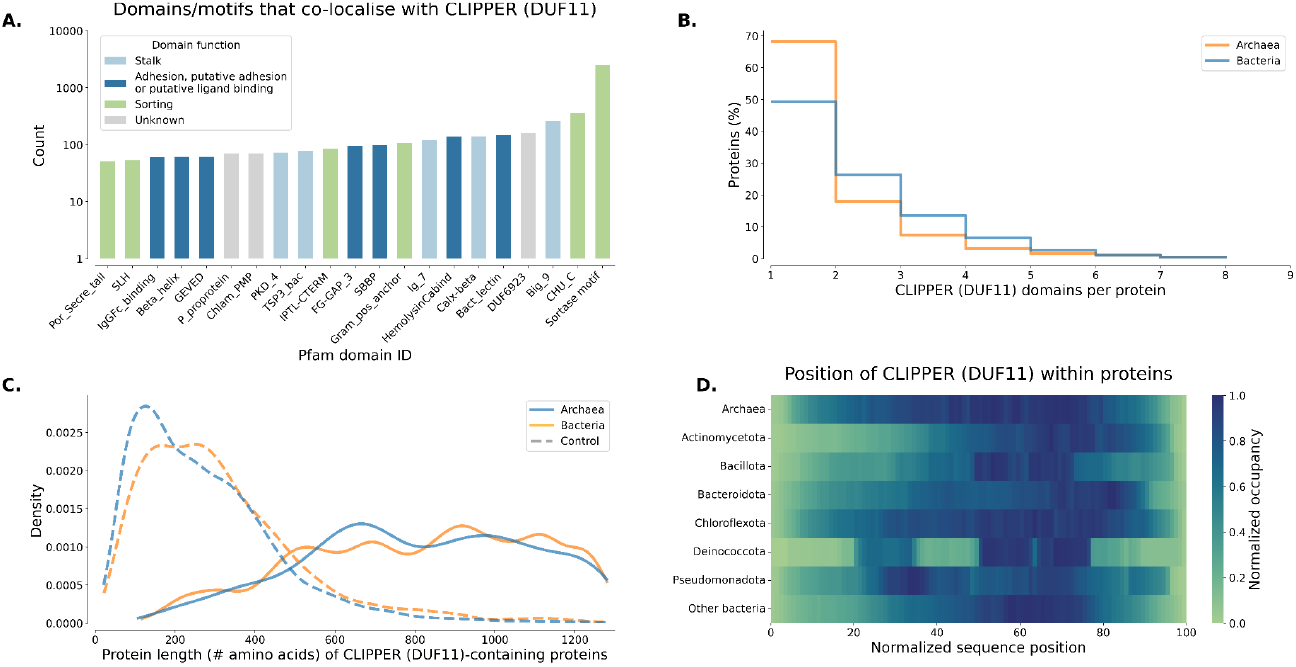
A. Pfam domains co-localising with isopeptide bond-containing CLIPPER domains in bacterial and archaeal AFDB proteins (domains detected <50 times are not shown). B. Histogram showing the number of isopeptide-containing CLIPPER repeats per AFDB protein. C. Protein length distribution of AFDB proteins with at least one isopeptide-containing CLIPPER domain, compared to control sequences as described in Figure 3. D. Positioning of isopeptide-containing CLIPPER domains within AFDB polypeptide sequences of archaea and various bacterial phyla.

While most DUF11 domains are found in proteins from Gram-positive and Gram-negative bacteria, some have been identified in multiple cell-surface proteins from archaea. Tandem DUF11 repeats are present in proteins that resemble fibrillar adhesins of the archaean genus *Methanothermobacter* (Sumikawa et al. 2019) and are present in porins (mostly without isopeptide bond signatures) that are proposed to form part of the archaeal S-layer structure (Doloman et al. 2024). DUF11-containing *Methanothermobacter* proteins are known to stabilise cell aggregates, and while the role of DUF11 domains in these cell proteins remains cryptic, they are suspected to be important for stabilisation of the protein (Sumikawa et al. 2019).

DUF11 is closely related to DUF7507, a new Pfam family built from isopeptide bond-containing domains identified by Isopeptor. DUF7507 domains are more compact than DUF11, typically consisting of seven β-strands (as judged by the AF2 predictions). Both DUF11 and DUF7507 domains are frequently found in tandem within surface polypeptides of bacteria and archaea, often alternating between the two families in tandem. Given the likely propensity of DUF11 and DUF7507 domains to harbour intramolecular isopeptide bonds, we have renamed and refer to them hereafter as CLIPPER and CLIPPER_2 domains (**C**ross-**L**inked **I**so**P**eptide **P**rotein in the **E**xtracellular **R**egion). Despite their wide phyletic distribution in a variety of putative host-binding fibrillar adhesins, neither CLIPPER nor CLIPPER_2 had previously been structurally or biophysically characterised. For this reason, we proceeded to determine the structure of a member of the CLIPPER family using X-ray crystallography to confirm the presence of an intramolecular isopeptide bond and undertook biophysical studies to characterise the thermal and proteolytic resilience of this domain.

### Structural and biophysical characterisation of a CLIPPER domain

We chose to structurally and biophysically characterise a CLIPPER domain from a putative fibrillar adhesin utilised by a Gram-negative bacterium within a well-characterised, high-quality genome. To this end, we identified a putative fibrillar adhesin 2129 amino acids in length from the genome of *Acinetobacter silvestris* ANC 4999 (Nemec et al. 2022), named B9T28_05395 (Uniprot ID: A0A1Y3CHT7_9GAMM). This fibrillar adhesin appears to be composed of an N-terminal adhesive CshA_NR2-like domain, a CshA_GEVED-like domain, a tandem repeat stalk of either CLIPPER domains or C-terminal cadherin-like domains, and an OmpA-like anchoring domain (Figure 5A). The CLIPPER repeat between residues 708 - 828 demonstrated the highest average identity to the other CLIPPER repeats (data not shown) and was subsequently chosen for experimental characterisation.

**Figure 5.**
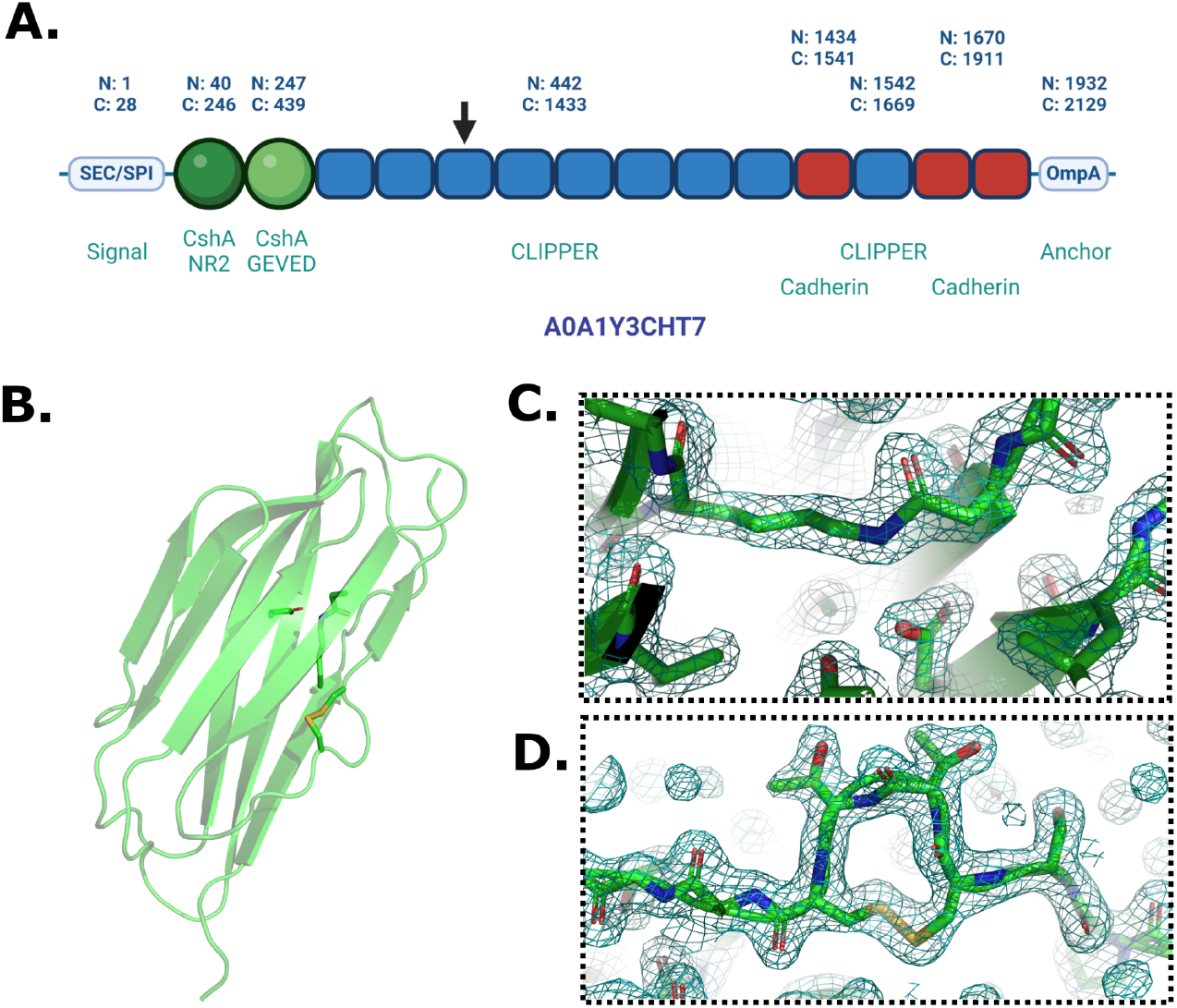
A. Schematic of fibrillar adhesin B9T28_05395 from *Acinetobacter silvestris* ANC 4999, with N- and C-termini of domain regions indicated. The black arrow indicates the CLIPPER domain investigated in this work (residues 708 - 828). B. A PyMOL rendering of the X-ray crystal structure of CLIPPER_WT_, depicted in cartoon format. C. A zoomed-in view of the isopeptide bond and putative catalytic aspartate of the CLIPPER_WT_ domain, depicted in stick format. D. A zoomed-in view of the CTTC disulphide motif. Electron density maps are shown in teal and contoured to 1σ (0.314 e/Å^3^).

Constructs of wild-type CLIPPER (CLIPPER_WT_) and a variant lacking the isopeptide-forming lysine (CLIPPER_K715A_) were expressed, purified (Figure S7), and cleaved of their His-tag prior to crystallisation attempts. Following sparse-matrix screening of various crystallisation conditions, CLIPPER_WT_ crystals were acquired, with data extending to 1.77 Å resolution (Table 2). The final modelled structure consists of a β-sandwich formed by two antiparallel sheets of four and five β-strands (Figure 5B). The domain presents as a CnaA-like fold and exhibits an intramolecular isopeptide bond between Lys-715 and Asn-806, presumably catalysed by nearby Asp-748 (Figure 5C). The final β-strand (β9) of the fold exhibits a mid-strand β-bulge that is caused by a tetrapeptide disulfide motif of Cys-818/Thr-819/Thr-820/Cys-821 (Figure 5D).

**Table 2.** Crystallography data collection and refinement statistics. Values in parentheses are for the highest resolution shell.

| Crystallography data collection and refinement |  |
| --- | --- |
| Data Collection |  |
| Space group | $P6_3$ |
| Unit cell dimensions (Å): A, B, C | 93.859, 93.859, 21.994 |
| Unit cell angles (°): $\alpha$, $\beta$, $\gamma$ | 90, 90, 120 |
| Crystal system | Hexagonal |
| Datasets merged | 2 |
| Resolution (Å) | 81.28–1.77 (1.80–1.77) |
| CC<sub>1/2</sub> (%) | 0.993 (0.401) |
| R<sub>pim</sub> | 0.161 (3.626) |
| Number of unique reflections | 11,209 (547) |
| Multiplicity | 38.7 (29.5) |
| Overall signal-to-noise ratio (I/$\sigma$) | 5.1 (0.4) |
| Completeness (%) | 99.91 (97.16) |
| Refinement |  |
| R<sub>work</sub> | 0.200 |
| R<sub>free</sub> | 0.209 |
| N<sub>o</sub> protein atoms used in refinement | 876 |
| N<sub>o</sub> water atoms used in refinement | 113 |
| B factors for protein atoms (Å<sup>2</sup>) | 31.89 |
| B factors for water atoms (Å<sup>2</sup>) | 36.29 |
| RMS deviations—length (Å) | 0.015 |
| RMS deviations—angle (°) | 1.544 |
| Ramachandran-favored residues (%) | 97 |
| Ramachandran outlying residues (%) | 1 |

We next sought to characterise the stabilising effects of the intramolecular isopeptide bond on CLIPPER. Since intramolecular isopeptide bonds usually impart significant thermotolerance to CnaA-like domains (Heidler et al. 2021), we investigated the thermal stability of CLIPPER_WT_ and the isopeptide-lacking CLIPPER_K715A_ using circular dichroism (CD) by collecting spectra from 5°C to 95°C at 5°C intervals (Figure 6A). CLIPPER_WT_ and CLIPPER_K715A_ demonstrate comparable β-sheet-like spectra, with a negative mean residue ellipticity (MRE) at 218 nm and positive MRE at <200 nm, indicating that both constructs are folded. At higher temperatures, a transition to a disordered state is observed for both constructs, at ∼80°C for CLIPPER_WT_ and ∼45°C for CLIPPER_K715A_ (Figure 6C). At high temperatures, the MRE values at 203 nm appear to differ between the polypeptides, reaching a plateau of ∼-7500 deg.cm^2^.dmol^-1^ in CLIPPER_WT_ and ∼-10,000 deg.cm^2^.dmol^-1^ in CLIPPER_K715A_ (Figures 6A and 6C), indicating that the intramolecular isopeptide bond may prevent total unfolding of the polypeptide chain.

**Figure 6.**
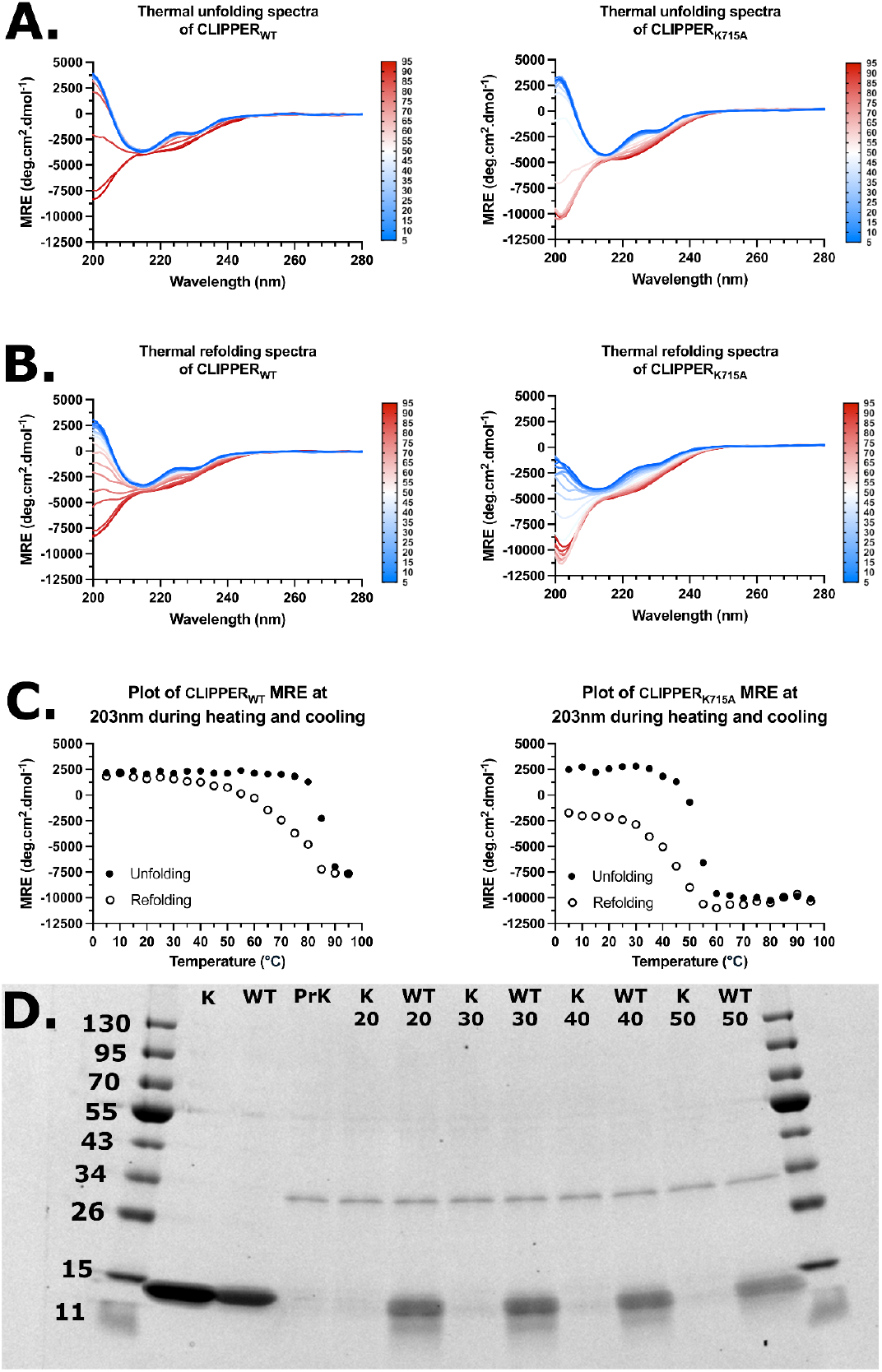
A. CD thermal unfolding spectra of CLIPPER_WT_ and CLIPPER_K715A_, increasing from 5°C to 95°C with spectra recorded at 5°C increments. B. CD thermal refolding spectra of CLIPPER_WT_ and CLIPPER_K715A_, decreasing from 95°C to 5°C with spectra recorded at 5°C increments. C. 203 nm MRE plot of CLIPPER_WT_ and CLIPPER_K715A_ during thermal unfolding and refolding. D. SDS-PAGE analysis of CLIPPER_WT_ (WT) and CLIPPER_K715A_ (K) polypeptides after incubation with buffer or Proteinase K (PrK) at the indicated temperatures for 1 hour.

We then assessed whether the intramolecular isopeptide bond facilitates refolding from thermally denatured states. To this end, the samples were subsequently cooled from 95°C to 5°C, with spectra collected at 5°C intervals (Figures 6B and C). During cooling, CLIPPER_WT_ appears to regain a predominantly β-sheet structure, whereas CLIPPER_K715A_ does not, with MRE between 200 - 205 nm failing to achieve a positive value. When plotting the folded-unfolded-refolded transition at 203 nm, both constructs demonstrate rapid unfolding but gradual refolding, which may be indicative of intermediate states during the refolding process (which is incomplete for CLIPPER_K715A_, Figure 6C). These data indicate that the intramolecular isopeptide bond of CLIPPER bestows significant thermotolerance to the domain and enables efficient refolding of the domain upon cooling, likely through intermediate states.

Finally, we sought to test the proteolytic susceptibility of CLIPPER_WT_ and CLIPPER_K715A_ to determine whether the intramolecular isopeptide bond imparts proteolytic resistance (as observed in other intramolecular isopeptide bond domains, Kang & Baker, 2009). Recombinant CLIPPER_WT_ and CLIPPER_K715A_ polypeptide were incubated with Proteinase K at 20°C, 30°C, 40°C and 50°C for 1 hour and visualised using sodium dodecyl sulphate polyacrylamide gel electrophoresis (SDS-PAGE). The results revealed that CLIPPER_WT_ is significantly more resistant than CLIPPER_K715A_ to proteolysis when treated with Proteinase K over all temperatures (Figure 6D), indicating that the presence of the isopeptide bond confers significant proteolytic resilience to the fold, in line with the properties of other intramolecular isopeptide bond domains (Kang & Baker, 2009).

## Discussion

This work characterised common environmental features surrounding intramolecular isopeptide bonds and surveyed their distribution in nature. Our analyses quantitatively confirm previous observations that these bonds are invariably found within hydrophobic cores, often near aromatic residues (Figure 2, Kang et al. 2007; Kang & Baker, 2009). These “aromatic caps” (as referred to herein) are detected in 71% and 41% of CnaA-like and CnaB-like domains respectively, in 86% and 24% of isopeptide bonds in *cis*/*trans* conformations, and are notably absent in folds containing Lys-Asp bonds. Additionally, water molecules are proximal to isopeptide Asn_Oδ_/Asp_Oδ_ atoms in more than half of the surveyed structures. Whether the presence or absence of water molecules and aromatic caps in structures harbouring intramolecular isopeptide bonds reflect distinct optimisations of the chemical environments remains unknown. It may be that the frequent occurrence of water molecules within the domain core represents water channels (as suggested previously, Hagan et al. 2010; Hu et al. 2011), or alternatively that they may play a stabilising role within the domain interior via key hydrogen-bond interactions. Additionally, considering their frequent presence, aromatic caps may play an important role in facilitation of isopeptide bond formation, or stabilisation of the fold more broadly. Further work may probe these features to elucidate their effect on isopeptide bond formation and domain stability.

Intramolecular isopeptide bond domains appear to group closely within three superfamily clans. Two extant domain families had not been annotated as intramolecular isopeptide bond-containing domain families prior to this work (SdrD-B and SpaA_3), while two (DUF11 and SpaA_3) were annotated as such in the Pfam database but lacked experimental validation. Ten new families were created, expanding the repertoire of known intramolecular isopeptide domain families to 26 (Table 1). Considering that all 26 families are predicted to resemble β-sandwich folds with Greek-key motifs, it is likely that CnaA-like and CnaB-like folds share a common evolutionary origin. Whether intramolecular isopeptide bonds are capable of forming in non-β-sandwich folds is not yet known, but recent work has introduced an autocatalytic intramolecular isopeptide bond to a β-sandwich fold lacking the Greek-key motif, indicating that such bonds can exist outside of CnaA/CnaB-like folds (Srisantitham et al. 2025). Future studies employing the findings of this work and protein design tools may elucidate whether isopeptide bonds can be engineered into alternative synthetic folds to generate stabilised synthetic domains for novel adhesive and material technologies.

Until now, only one intramolecular isopeptide bond domain has been structurally characterised from an organism that is not a Gram-positive bacterium (Heidler et al. 2021), and their wider phyletic distribution has remained uncertain (Schwarz-Linek & Banfield, 2014). We found that intramolecular isopeptide bonds are prevalent in Gram-positive bacteria, but that three families demonstrate wider distribution. DUF11 (renamed CLIPPER), and the closely related domains DUF7507 (renamed CLIPPER_2) and DUF7619 are frequently found in polypeptides of Gram-negative bacteria, while CLIPPER and CLPPER_2 also occur in archaeal proteins (Figure S3). We found that tandemly repeating intramolecular isopeptide bond domains are usually located in elongated cell-surface proteins of host-binding pathogens, opportunistic pathogens, and commensal microbes (Figures 3, S4 and Table S3). Notably, CLIPPER appears to be the most widely distributed domain, often tandemly repeated in the stalks of fibrillar adhesins (Figures 4 and S3). The frequent use of tandemly repeating intramolecular isopeptide bonds in fibrillar adhesins of host-binding bacteria and archaea indicates that they may be important for the efficient colonisation of host tissues and may, therefore, present attractive targets for novel antimicrobial therapeutics. Future work may focus on generating novel therapeutics to interfere with isopeptide bond formation and abrogate host colonisation by pathogens, following previous work that has demonstrated promising results employing this strategy (Rivas-Pardo et al. 2018).

Given that CLIPPER was found to be the most broadly distributed intramolecular isopeptide domain from our work, we experimentally characterised an exemplary domain from the fibrillar adhesin B9T28_05395 of *Acinetobacter silvestris* ANC 4999. Our crystal structure reveals that the CnaA-like fold harbours an intramolecular isopeptide bond between Lys-715 and Asn-806, presumably catalysed by Asp-748 (Figure 5). The isopeptide bond enables significant thermostability and resistance to proteolysis and can refold after thermal denaturation, properties that were abrogated in an isopeptide-lacking variant (Figure 6). This indicates that the isopeptide bond of CLIPPER domains enables significant resilience to stress, which is likely of functional importance when present in the stalks of fibrillar adhesins from Gram-positive bacteria, Gram-negative bacteria, and archaea. Whether other intramolecular covalent cross-link strategies exist within surface proteins of diverse phyla remains unknown. Follow-up studies investigating the distribution of other intramolecular covalent bonds (such as ester and thioester bonds, Walden et al. 2015) in the AFDB may reveal the phyletic diversity of such domains, the diversity among their families, and key features of their chemical environments.

## Materials and methods

### Analysis of intramolecular isopeptide bond structural features

Wild-type PDB structures containing an intramolecular isopeptide bond, with X-ray diffraction resolution ≤2.5 Å and no unusual residue properties (e.g. incomplete or missing side chains present on flexible loops), were chosen from the previously collated collection of intramolecular isopeptide bond domains (Costa et al. 2025). This dataset includes remodelled structures of intramolecular isopeptide bonds, which have been deposited in the PDB with incorrect isopeptide bonds geometries. Only one PDB entry was assessed per sequence-identical domain, considering the 20 residues flanking each side of the first and last isopeptide bond signature positions. rASA was calculated using the Biotite package v1.0.1 (Kunzmann et al. 2018) sasa function with point_number 500 and values normalised using Rost and Sander maximum ASA values (Rost and Sander, 1994). Bonds were classified as *cis* for pseudo ω angles <60° or *trans* for angles > 120°. “Aromatic caps” were identified as aromatic residues with the centroid of their aromatic ring within 6 Å of the isopeptide bond Lys_Nζ_ atom, where the angle between the centroid-Lys_Nζ_ atom vector was <30° or >150° from the normal of the aromatic ring plane (in the case of tryptophan, the closest of the two rings was considered). Where no aromatic caps were detected, the closest aromatic within 10 Å of the Lys_Nζ_ atom was plotted. The analyses were conducted with custom python v3.12.2 scripts with the Biotite, Pandas v2.1.1 (McKinney, 2010), seaborn v0.13.2 (Waskom, 2021), matplotlib v3.8.0 (Hunter et al. 2007), biopython v1.83 (Cock et al. 2009) and Numpy v1.26.4 (Harris et al. 2020) packages for calculations.

### AlphaFold2 modelling

AlphaFold2 modelling was performed on A100 GPUs with AlphaFold version 2.3.1 (Jumper et al. 2021) (https://github.com/kalininalab/alphafold_non_docker) and cuda version 11, with active amber relaxation and PDB templates options. Multiple sequence alignment (MSA) was performed on the sequence database suggested from the GitHub webpage on 07/2021 (https://github.com/kalininalab/alphafold_non_docker). Missing residues from PDB sequences were replaced with glycine for AF2 predictions. Predictions generated 5 models, and the highest pLDDT-scoring model was selected.

### Large-scale prediction of intramolecular isopeptide bonds using Isopeptor, Pfam mapping and creation of new Pfam domains

Isopeptor version v0.0.75 was installed using the pip python package manager and run with default parameters, employing a probability threshold of >0.65 against the AFDB database version 4 and the BFVD database version 2023_02. AFDB domains detected by Isopeptor were mapped to Pfam version 37_0 entries and assigned to Pfam domains when entries covered at least two of the three residues required for intramolecular isopeptide bond formation. Sortase motifs were annotated within isopeptide bond-containing proteins using the following regex expressions within the last 50 C-terminus amino acids: LP.T[G|A|N|D], NP.TG, LP.GA, LA.TG, NPQTN, IP.TG. Newly created pfam domains were mapped to AFDB proteins using hmmscan (HMMER version 3.3.2, Eddy, 2011) and filtered considering annotated domain and sequence gathering thresholds. In cases of conflicting annotations, priority was given to domains annotated in Pfam version 37_0. AFDB hits with no corresponding Pfam annotation were used to create new Pfam domain families via sequence clustering, using N-terminal domain boundaries -10/-5 residues from the isopeptide-bonded lysine, and C-terminal domain boundaries +5/+30 residues from the isopeptide-bonded asparagine/aspartate for CnaB-like/CnaA-like domains, respectively. Clustering was performed using MMseqs2 software version 17.b804f (Steinegger & Soding, 2017) via the easy-cluster command and the following flags: cluster-mode 2, cov-mode 0, min-seq-id 0.25, coverage 0.9, and aligned using MAFFT or Muscle (Edgar 2004; Katoh & Standley, 2013). The resultant MSAs were used as initial seeds to search the reference proteome database using hmmsearch (HMMER version 3.3.2). Pfam families were built via repeated iterative searches and manual refinement of boundaries, member selection and inclusion thresholds (which were adjusted for each family to exclude false positives and optimise signal-to-noise).

### Assessing fibrillar adhesin prevalence

For the prediction of fibrillar adhesin prevalence, the FAL_prediction software was downloaded and employed (https://github.com/VivianMonzon/FAL_prediction). FAL_prediction was employed against full-length sequences of Isopeptor-identified AFDB proteins, using a probability threshold of >0.9. The Iupred2a and T-Reks dependencies were downloaded from https://iupred2a.elte.hu/download_new (Mészáros, Erdős & Dosztányi, 2018) and https://bioinfo.crbm.cnrs.fr/index.php?route=tools&tool=3 (Jorda & Kajava, 2009), respectively.

### CLIPPER expression

A synthetic gene encoding residues 708 – 828 of *Actinetobacter silvestris* ANC4999 fibrillar adhesin B9T28_05395 (Uniprot accession A0A1Y3CHT7, “CLIPPER_WT_”) and a mutant lacking the isopeptide-forming lysine residue (“CLIPPER_K715A_”) were ordered from Thermo Fisher Scientific (Table S4), codon-optimized for *E. coli*. Synthetic genes were inserted into HindIII/KpnI-linearised pOPINF expression vector following polymerase chain reaction with appropriate primers, using the In-Fusion™ kit (Takara Bio), following the instructions of Berrow *et al.* 2007 and verified via DNA sequencing (Eurofins Genomics). The encoded constructs contain an N-terminal hexa-histidine tag with a 3C cleavage site, and were transformed into BL21(DE3) *Escherichia coli* (Thermo Fisher Scientific) using heat-shock, grown in 1 L cultures of Luria Broth supplemented with carbenicillin at 100 μg.mL^-1^ until an optical density of 0.8, and induced via addition of isopropyl β-D-1-thiogalactopyranoside to a concentration of 1 mM. Cultures were incubated for 16 hours at 18°C (180 RPM), harvested by centrifugation (4,500 x g, 30 minutes), the pellets flash-frozen in liquid nitrogen, and stored at -80°C.

### CLIPPER purification

Pellets were resuspended in phosphate-buffered saline (PBS, 137mM NaCl, 3mM KCl, 10mM Na_2_HPO_4_, 1.8mM KH_2_PO_4_, pH 7.4), lysed using a 120 sonic dismembrator (Fisher Scientific, amplitude 70%, 20 second on, 20 second off, 10 minutes) and centrifuged (39,000 x g, 30 minutes). Supernatant was applied to a 5 mL HisTrap Nickel-NTA column (Cytiva) equilibrated with PBS + 20mM Imidazole, pH 7.4. A concentration gradient of 20 mM to 500 mM imidazole in PBS was applied over 60 mL for 1 hour. Fractions were collected, concentrated and subjected to size exclusion chromatography (SEC) purification using an EnRich SEC 650 10 × 300 column (BIORAD) equilibrated with PBS for circular dichroism and proteolysis assays or Tris-buffered saline (TBS, 20 mM Tris, 100 mM NaCl, pH 8.0) for crystallisation studies. SEC fractions were subjected to SDS-PAGE analysis, mixing 10 μL with 10 μL of 2x Laemli buffer (BioRad), heated at 95°C for five minutes, applied to Novex Tris-Glycine precast SDS-PAGE gels (Thermo), and subjected to 200 V for 30 minutes in an X-Cell SureLock system (Thermo), and visualised using Instant Blue stain (Fisher Scientific).

### CLIPPER crystallisation and data collection

His-tags of CLIPPER_WT_/CLIPPER_K517A_ were cleaved using HRV3C protease (Takara Bio), removed via application to a HisTrap column, and repurified using SEC. Sitting-drop vapour-diffusion sparse matrix screens of CLIPPER_WT_/CLIPPER_K517A_ at 5 – 10 mg.mL^-1^ were set up using commercial screens (Molecular Dimensions). Well-diffracting crystals were grown in 1:1 droplet ratios of 5 mg.mL^-1^ CLIPPER_WT_ with 0.2 M Ammonium Sulphate, 0.2 M Sodium/Potassium Tartrate, pH 5.5 in 4 μL droplets (CLIPPER_K715A_ yielded no crystals). Crystals were mounted in LithoLoops (Molecular Dimensions), flash-cooled in liquid nitrogen without cryoprotectant, and diffraction data collected at the I24 beamline of Diamond Light Source, UK, on the i24 beamline. Data was processed using Xia2 and DIALS (Winter, 2010; Winter et al. 2018) and phases calculated using molecular replacement in CCP4i2 with MOLREP (Vagin & Teplyakov, 1997; Potterton et al. 2018) using the CLIPPER_WT_ AF2 model, and iteratively refined via manual model building with REFMAC and COOT (Vagin et al. 2004; Emsley et al. 2010). The model and structure factors for CLIPPER_WT_ have been deposited at the PDB with the code 9IFR.

### Circular dichroism

1 mL aliquots of CLIPPER_WT_/CLIPPER_K517A_ polypeptides were dialysed overnight at 4°C in 5 L of circular dichroism (CD) buffer (10 mM sodium phosphate, 100 mM sodium fluoride, pH 7.4) using SnakeSkin dialysis tubing (10 kDa cutoff, Thermo). CD spectra were recorded for CLIPPER_WT_ and CLIPPER_K517A_ using a Jasco J-1500 spectrophotometer continuously purged with nitrogen and fitted with a Peltier temperature control unit. Spectra were obtained from sample volumes of 300 μL in a cuvette with a 1 mm path length (Hellma Analytics) with protein concentrations of 0.36 mg.mL^-1^ (CLIPPER_WT_) and 0.38 mgmL^-1^ (CLIPPER_K715A_) after blanking with CD buffer from the dialysis bucket. Spectra were recorded from 5 – 95 °C at 5 °C intervals (thermal unfolding) and subsequently 95 – 5 °C (thermal refolding) over a spectral range of 180 – 280 nm. High-tension threshold (HT) voltage was continuously recorded to ensure HT was <600 V. Spectra were averaged from four repeat scans and smoothed using a Savitsky–Golay smoothing algorithm over a window of 11 data points.

### Proteolysis assay

Proteinase K from *Tritirachium album* was used for the proteolysis assay (Sigma Aldrich). 10 μL of 1 mgmL^-1^ CLIPPER_WT_/CLIPPER_K517A_ were mixed with 10 μL of 1 mg.mL^-1^ Proteinase K in PBS and aliquoted into thin-walled PCR tubes (StarLab) on ice. After mixing, aliquots were placed inside of a Sensoquest LabCycler (Geneflow) along a heated gradient at 20°C, 30°C, 40°C or 50°C for 1 hour. Aliquots were immediately mixed with SDS running buffer and subjected to SDS-PAGE analysis.

## Author contributions

**Francesco Costa** – Conceptualisation; Data Curation; Formal Analysis; Investigation; Methodology; Software; Validation; Visualisation; Writing – original draft; Writing – review and editing. **Ioannis Riziotis** - Conceptualisation; Methodology; Software; Validation. **Antonina Andreeva** - Conceptualisation; Data Curation; Formal Analysis; Investigation; Methodology; Visualisation; Writing – original draft; Writing – review and editing. **Delhi Kalwan** – Investigation; Writing – review and editing. **Jennifer de Jong** - Investigation; Writing – review and editing. **Philip Hinchliffe** – Formal analysis; Validation; Writing – review and editing. **Fabio Parmeggiani** - Funding acquisition; Resources; Supervision; Writing – review & editing. **Paul Race** – Conceptualisation; Funding acquisition; Resources; Supervision; Writing – review & editing. **Steve Burston** - Conceptualisation; Methodology; Supervision; Writing – review & editing. **Alex Bateman** - Conceptualisation; Funding acquisition; Methodology; Project administration; Writing – original draft; Writing – review and editing. **Rob Barringer** - Conceptualisation; Data Curation; Funding acquisition; Investigation; Methodology; Project administration; Supervision; Visualisation; Writing – original draft; Writing – review and editing.

## Funding sources

This work was supported by the UKRI Biotechnology and Biological Sciences Research Council [BB/X012492/1, BB/W013959/1, BB/T008741/1 and BB/T001968/1], EMBL core funds, and a UKRI Engineering and Physical Sciences Research Council Impact Accelerator Award [EP/X525674/1].

## Acknowledgments

The authors would like to thank Diamond Light Source for beamtime (proposal MX31440) and the staff of beamline i24 for their assistance. We would also like to thank Angela Nobbs for discussions regarding fibrillar adhesins and mechanisms of colonisation, Alexandr Nemec for discussions regarding genetic diversity of *Acinetobacter*, and Uli Schwartz-Linek for discussions regarding intramolecular isopeptide bond formation, and associated properties. We also thank Thom Sharp for hosting Rob Barringer during the period of this work, and Ross Anderson for allowing the use of a Jasco J-1500 spectrophotometer for circular dichroism studies.

## Data availability statement

The data that support the findings of this study are openly available in Zenodo at http://doi.org/10.5281/zenodo.15024939. The code used for the analysis is available at: https://github.com/FranceCosta/isopeptide_bonds_global_survey.

## Conflict of Interest statement

The authors declare no conflicts of interest.

## Notes

https://zenodo.org/records/15024939

https://github.com/FranceCosta/isopeptide_bonds_global_survey

### Competing Interest Statement

The authors have declared no competing interest.

