## Supplementary information for "A global survey of intramolecular isopeptide bonds"

### Supplementary tables

| Intramolecular isopeptide bond type | Intramolecular isopeptide bond-containing folds with a water molecule within 5 Å from the bond Oδ |
|---|---|
| **CnaA-like** | 59% |
| **CnaB-like** | 54% |

**Table S1:** Prevalence of proximal waters and oxygen atoms <5 Å of intramolecular isopeptide bonds. Only one PDB entry was assessed per sequence-identical domain. “Non-catalytic oxygen” is any oxygen that does not belong to the isopeptide bond catalytic residue or nearby water molecules.

| Pfam ID (accession code) | Pfam Clan ID (accession code)-isopeptide bond class type | Total number of domains present in the AFDB | Predicted to contain an intramolecular isopeptide bond |
|---|---|---|---|
| Collagen_bind (PF05737) | Adhesin (CL0204) CnaA-like | 2,589 | 2,307 |
| GramPos_pilinBB (PF16569) | | 2,226 | 2,044 |
| AgI_II_C2 (PF17998) | | 876 | 684 |
| Antigen_C (PF16364) | | 316 | 246 |
| Sgo0707_N2 (PF20623) | | 274 | 171 |
| DUF7926 (PF25548) | | 181 | 148 |
| DUF7929 (PF25551) | | 162 | 123 |
| DUF7925 (PF25546) | | 154 | 109 |
| DUF11 (PF01345) | E-set (CL0159) CnaA-like | 25,913 | 11,756 |
| DUF7507 (PF24346) | | 5,318 | 3,146 |
| DUF7619 (PF24595) | | 1,981 | 1,000 |
| DUF7933 (PF25564) | | 776 | 247 |
| GBS104-like_Ig (PF21426) | | 425 | 235 |
| DUF7927 (PF25549) | | 524 | 231 |
| DUF7617 (PF24593) | | 78 | 73 |
| SpaA (PF17802) | Transthyretin (CL0287) CnaB-like | 27,202 | 16,929 |
| Cna_B (PF05738) | | 6,407 | 5,013 |
| FctA (PF12892) | | 3,002 | 2,170 |
| DUF5979 (PF19407) | | 2,528 | 1,965 |
| GramPos_pilinD1 (PF16555) | | 2,784 | 1,692 |
| DUF7601 (PF24547) | | 818 | 668 |
| SpaA_4 (PF24514) | | 567 | 378 |
| SpaA_2 (PF19403) | | 1,259 | 246 |
| SpaA_3 (PF20674) | | 622 | 196 |
| GramPos_pilinD3 (PF16570) | | 165 | 137 |
| SdrD_B (PF17210) | | 9,185 | 36 |

**Table S2**: Pfam domains detected with Isopeptor and total counts of domains from the AFDB. Domain assignment was executed as explained in the results section. Only domains detected at least 20 times are shown. False positives have been excluded.

| Gram-positive pathogens | Gram-negative pathogens |
|---|---|
| Abiotrophia defectiva | Acinetobacter baumannii |
| Actinomyces naeslundii | Acinetobacter haemolyticus |
| Actinomyces oris | Brucella abortus |
| Bacillus anthracis | Brucella melitensis |
| Bacillus cereus | Brucella suis |
| Bacillus thuringiensis | Catonella morbi |
| Clostridioides difficile | Legionella bozemanae |
| Clostridium botulinum | Porphyromonas gingivalis |
| Clostridium perfringens | Salmonella enterica |
| Clostridium tetani | Salmonella newport |
| Hathewaya histolytica (Clostridium histolyticum) | Salmonella typhimurium |
| Corynebacterium diphtheriae | Vibrio alginolyticus |
| Corynebacterium jeikeium | Vibrio parahaemolyticus |
| Corynebacterium striatum | |
| Enterococcus faecalis | |
| Enterococcus faecium | |
| Listeria grayi | |
| Listeria ivanovii | |
| Listeria monocytogenes | |
| Staphylococcus aureus | |
| Streptococcus agalactiae | |
| Streptococcus anginosus | |
| Streptococcus downei | |
| Streptococcus dysgalactiae | |
| Streptococcus gordonii | |
| Streptococcus mitis | |
| Streptococcus mutans | |
| Streptococcus oralis | |
| Streptococcus pneumoniae | |
| Streptococcus pyogenes | |
| Streptococcus sanguinis | |

**Table S3:** A list of human-binding pathogens/opportunistic pathogens that employ cell-surface proteins containing intramolecular isopeptide bonds identified by Isopeptor.

| | |
|---|---|
| **Gene constructs (codon optimised for Escherichia coli)** | |
| **CLIPPER$_{WT}$** | AAACCGACCGTTGATGTTGTTAAAACCACCACCGCAACCACCGCCAAAGTTGGTG<br>ATACCATTGATTATACCGTTAAAGTTACCGTTGCAAATAGCCAGACCACCGATGC<br>ACTGACCCTGAATGATACCTTAGGTCAGGGTCTGAGCTTTGTTAGCGGCACCGCA<br>CCGACCGGTTGGACCCTGACAGGTAATGGTCAGGCAATTAACATTGCAGCACCGA<br>AAGGTCAGATTCCGGGTACATATAATCTGACGTATAAAGTTCTGGTTGGCACCGA<br>TGCCGTTAATAATGTTGTGAATAAAGTGACCGCAAGCGGTGGTGATAAACCGAGC<br>TGCACCACCTGTACCACCACCACACCGGTTACC |
| **CLIPPER$_{K715A}$** | AAACCGACCGTTGATGTTGTTGCAACCACCACCGCAACCACCGCCAAAGTTGGTG<br>ATACCATTGATTATACCGTTAAAGTTACCGTTGCAAATAGCCAGACCACCGATGC<br>ACTGACCCTGAATGATACCTTAGGTCAGGGTCTGAGCTTTGTTAGCGGCACCGCA<br>CCGACCGGTTGGACCCTGACAGGTAATGGTCAGGCAATTAACATTGCAGCACCGA<br>AAGGTCAGATTCCGGGTACATATAATCTGACGTATAAAGTTCTGGTTGGCACCGA<br>TGCCGTTAATAATGTTGTTAATAAAGTGACCGCAAGCGGTGGTGATAAACCGAGC<br>TGCACCACCTGTACCACCACCACACCGGTTACC |
| **Primers used in this study** | |
| **CLIPPER constructs Forward primer** | **<u>AAGTTCTGTTTCAGGGCCCG</u>**AAACCGACCGTTGATGTTGTT |
| **CLIPPER constructs Reverse primer** | **<u>ATGGTCTAGAAAGCTTTA</u>**GGTAACCGGTGTGGTGGTG |
| **Plasmid used in this study** | |
| **pOPINF** | Ampicillin-resistant expression vector for recombinant proteins, incorporating an N-terminal cleavable hexa-histidine tag (MAHHHHHHSSGLEVLFQGP, Berrow et al. 2007) |

**Table S4:** Sequences of synthetic DNA used in this study, and the source of pOPINF plasmid used for protein expression. pOPINF-complementary sequences of primers are underlined in bold.

### Supplementary figures

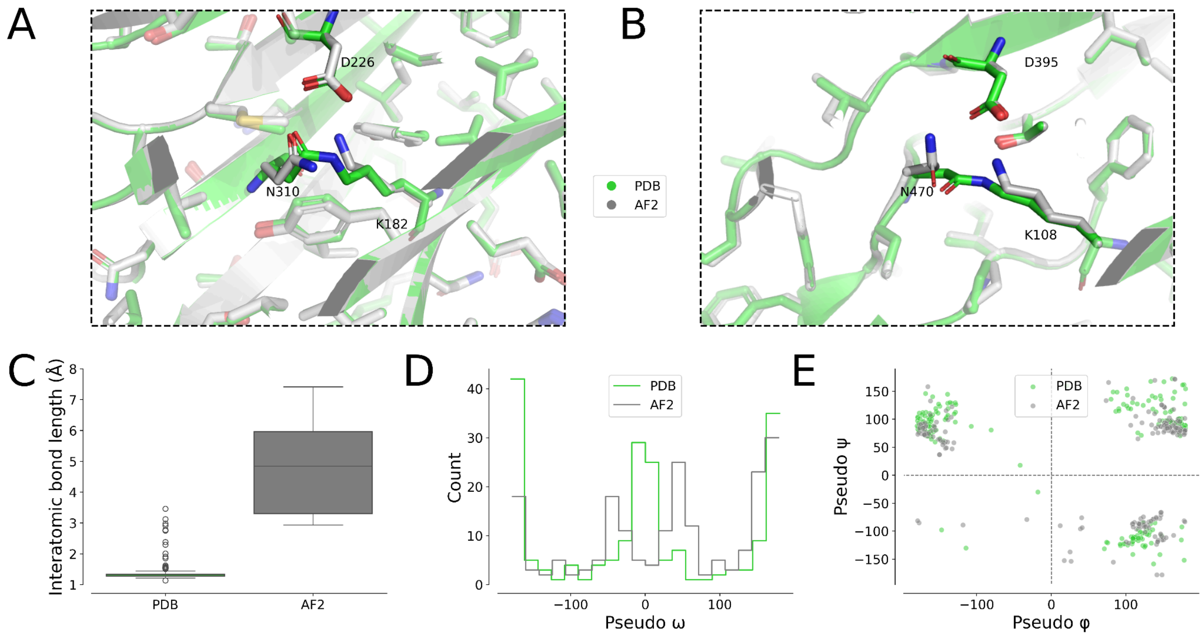


**Figure S1**: A. An example in which the intramolecular isopeptide bond modelled by AlphaFold2 resembles the bond of the PDB structure (PDB ID: 5XCB). B. An example in which the intramolecular isopeptide bond modelled by AlphaFold2 does not resemble the one found in the PDB structure (PDB ID: 6M3Y). For panels A and B, both intramolecular isopeptide bonds are confidently predicted by Isopeptor (probability 0.89 and 0.73, respectively). The average pLDDT of isopeptide bond side chains in the AlphaFold2 models is >90. C. Isopeptide bond length distributions as calculated between the atoms that form the covalent link ($Lys_{N\zeta}$ and $Asp/Asn_{C\gamma}$). D. Dihedral pseudo ω angle distributions. E. pseudo ψ and φ angle distributions. Pseudo dihedral angles were calculated as described in Costa et al. 2025. The difference in distances between asparagine and lysine side chains in AF2 models likely reflects an attempt to prevent local clashes. This is also reflected in the distribution of pseudo ω angles (in which the *cis* conformation is unfavourable in AlphaFold2 models).

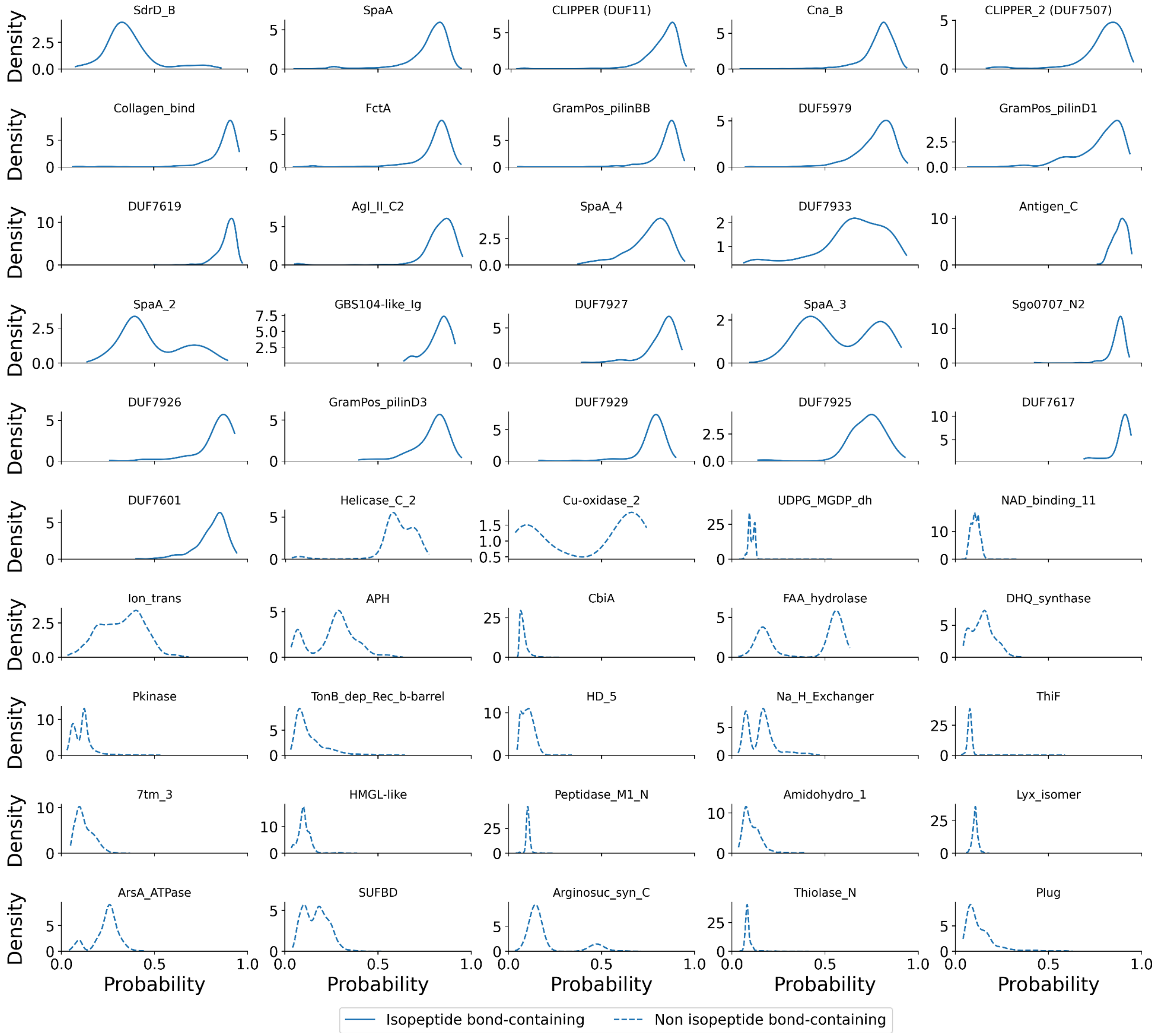


**Figure S2**: Distribution of Isopeptor probabilities for domains which contain an intramolecular isopeptide bond and for domains unlikely to contain an isopeptide bond. Intramolecular isopeptide bond-containing domains exhibit a higher Isopeptor probability score (above the threshold of 0.65), indicating that Isopeptor is detecting isopeptide bonds with high recall. There are a few exceptions to this: SpaA_2, SpaA_3 and SdrD_B domains have a substantial fraction of isopeptide bonds predicted with low confidence, likely due to a lack of experimental templates closely matching their isopeptide bond geometries. Helicase_C_2 and Cu-oxidase_2 domains, which have a substantial fraction of isopeptide bond signatures above the probability threshold of 0.65, were determined to be false positives.

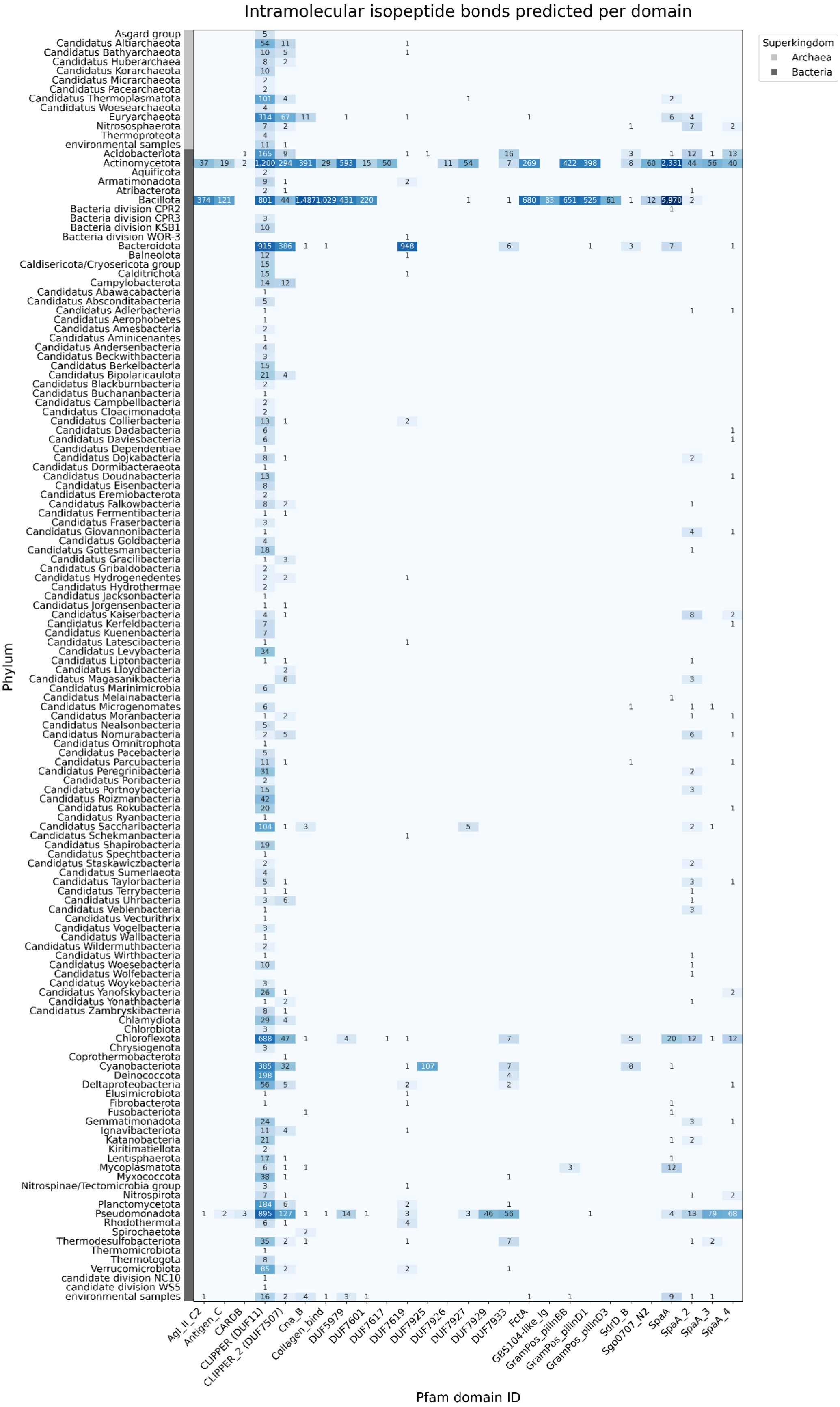


**Figure S3**: Distribution of intramolecular isopeptide bond-containing domains per phylum and superkingdom, detected with Isopeptor.

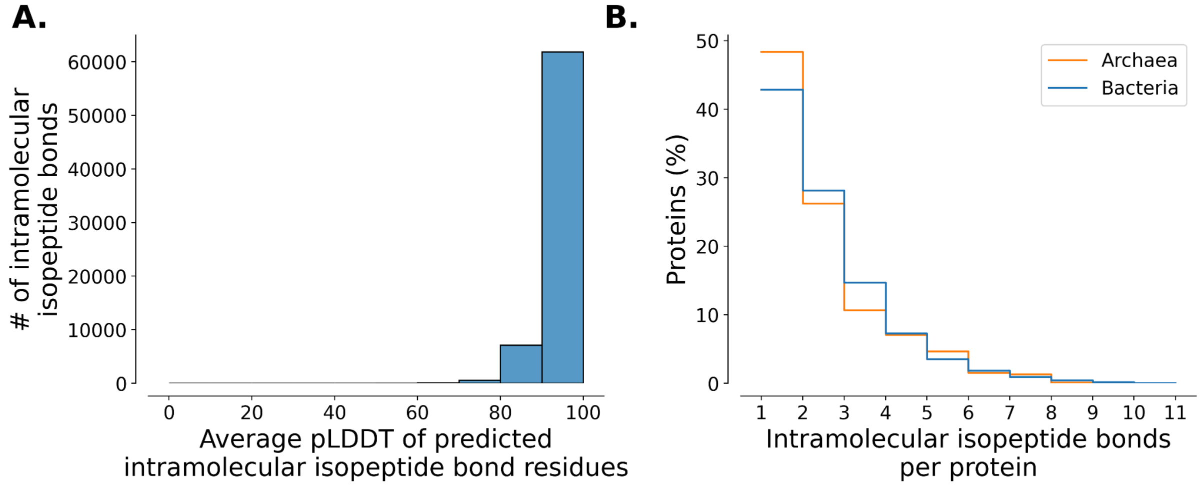


**Figure S4**: A. Distribution of pLDDT values for predicted intramolecular isopeptide bond residues across the AFDB. B. Intramolecular isopeptide bonds detected per protein. 57% and 52% of bacterial and archaeal proteins detected by Isopeptor contain more than one isopeptide bond, respectively.

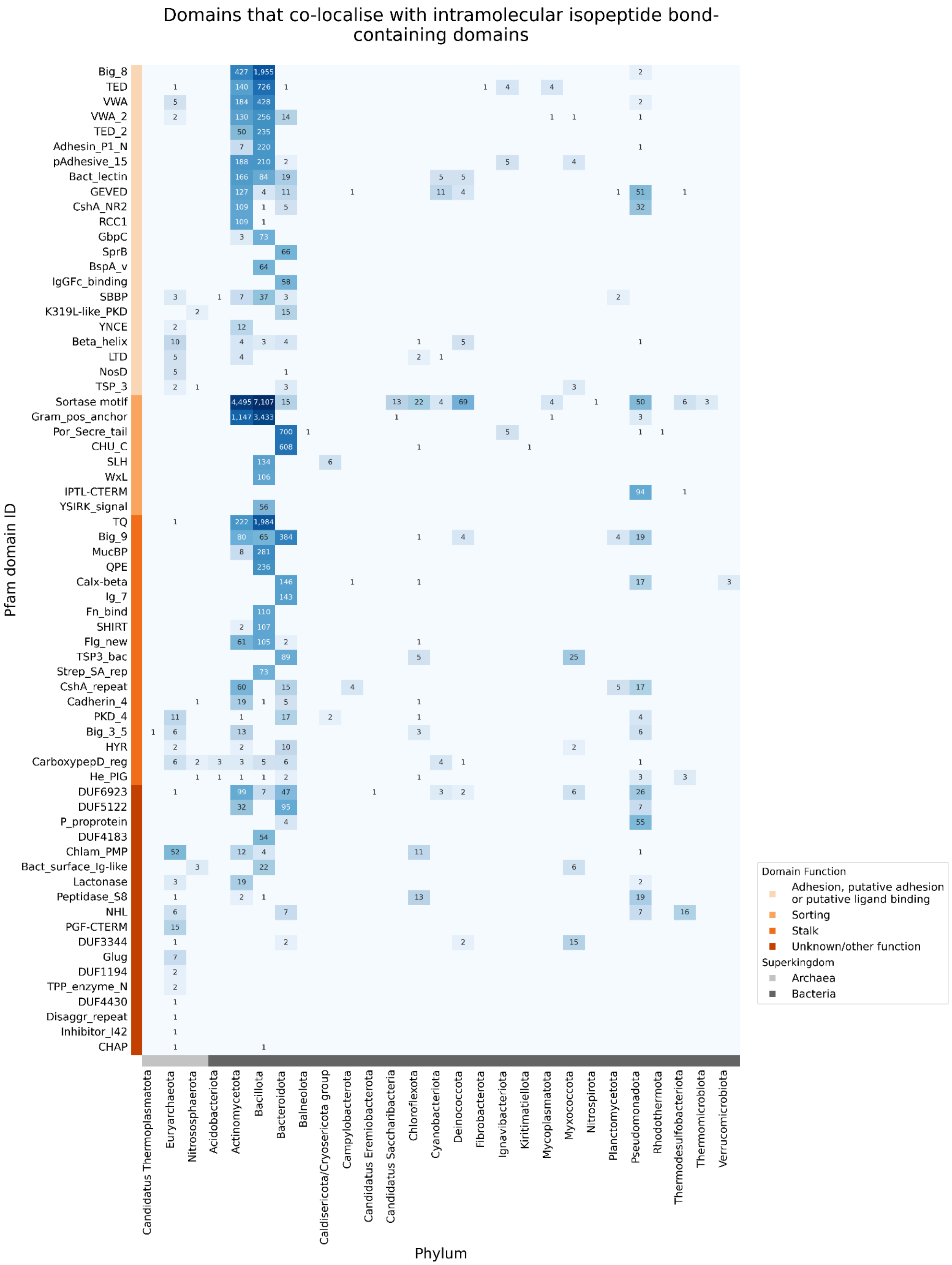


**Figure S5:** Domains neighbouring intramolecular isopeptide bond-containing domains, divided into functional classes and phylum. The most common functional classes are adhesion, sorting and stalk.

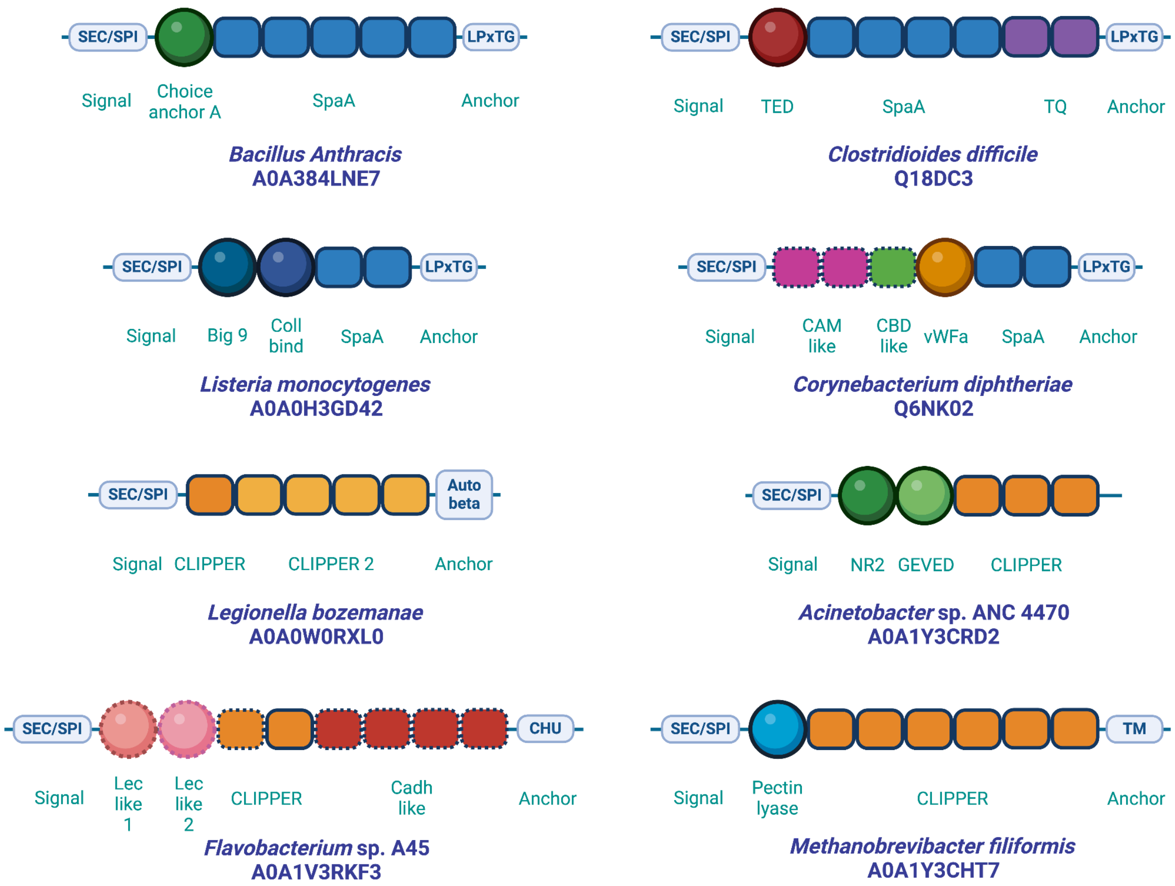


**Figure S6.** A selection of AFDB proteins predicted to contain intramolecular isopeptide domains, identified by Isopeptor. Adhesive domains are depicted as circles, stalk domains are depicted as rounded boxes, and anchor motifs/domains and signal peptides are depicted in light blue boxes at the N and C termini, respectively. Domains are colour coded by domain identity. Dotted outlines indicate domains that could not be correlated to Pfam domain families using sequence information, but that exhibit predicted structures that are similar to other domains of known function. Domain families are listed below each domain or domain repeat region. TED = Thioester Domain, TQ = T-Q ester bond domain, Coll bind = Collagen binding domain, CAM like = Cell adhesion module-like domain, CBD like = Carbohydrate binding-like domain, Lec like = Lectin-like domain, Cadh like = Cadherin-like domain.

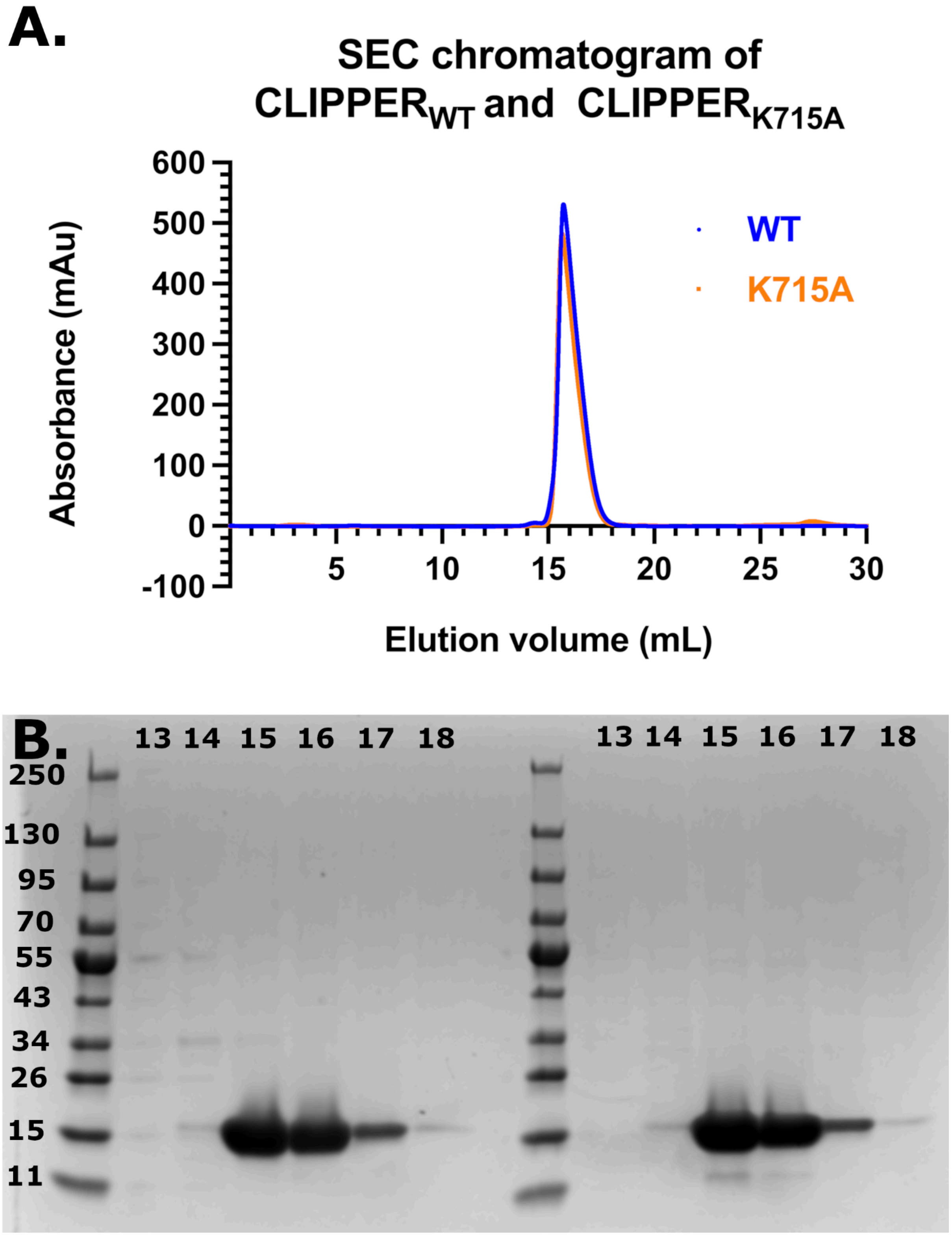


**Figure S7.** A. SEC chromatograms of $CLIPPER_{WT}$ and $CLIPPER_{K715A}$, demonstrating one primary peak per polypeptide indicative of a monomer. B. SDS-PAGE analysis of eluted SEC fractions of $CLIPPER_{WT}$ (left) and $CLIPPER_{K715A}$ (right).
